# Identifying proximal RNA interactions from cDNA-encoded crosslinks with ShapeJumper

**DOI:** 10.1101/2021.06.10.447916

**Authors:** Thomas W. Christy, Catherine A. Giannetti, Alain Laederach, Kevin M. Weeks

## Abstract

SHAPE-JuMP is a concise strategy for identifying close-in-space interactions in RNA molecules. Nucleotides in close three-dimensional proximity are crosslinked with a bi-reactive reagent that covalently links the 2’-hydroxyl groups of the ribose moieties. The identities of crosslinked nucleotides are determined using an engineered reverse transcriptase that jumps across crosslinked sites, resulting in a deletion in the cDNA that is detected using massively parallel sequencing. Here we introduce ShapeJumper, a bioinformatics pipeline to process SHAPE-JuMP sequencing data and to accurately identify through-space interactions. ShapeJumper identifies proximal interactions with near-nucleotide resolution using an alignment strategy that is optimized to tolerate the unique non-templated reverse-transcription profile of the engineered crosslink-traversing reverse-transcriptase. JuMP-inspired strategies are now poised to replace adapter-ligation for detecting RNA-RNA interactions in most crosslinking experiments.

## Introduction

RNA molecules form multiple levels of intra- and inter-molecular higher order structure, and these structures often have important functions. Secondary structures form via base pairing, and secondary structures may further fold into compact tertiary structures mediated by interactions involving canonically or non-canonically paired nucleotides^1,2^. Developing robust models of RNA secondary and tertiary structure is an important first step in understanding the underlying function of an RNA, and defining well-determined structures can lead to identification of novel functional elements^3,4^. Notable progress has been made using chemical probing experiments to broadly and accurately map biologically relevant secondary structures^5–7^. In contrast, efficient experimental mapping tertiary interactions remains a challenging, unresolved problem^8,9^, although notable progress is being made^10–12^.

In principle, RNA crosslinking should be able to identify short through-space interactions. Chemical probes such as psoralen analogs^13–16^, formaldehyde^17,18^ and bis-succinimidyl esters^18^, and short wavelength ultraviolet (UV) irradiation^18–20^ have been used to crosslink interacting nucleotides. In practice, identifying the precise locations of RNA crosslinks is difficult^8,9,21^. Recent, potentially high-throughput, methods to read out RNA-RNA crosslinks rely on variants of proximity ligation to identify crosslinked nucleotides^13–20^. Typically, RNAs are crosslinked and then some combination of RNA fragmentation, crosslink capture, and enrichment is used to obtain linked RNAs whose ends are close to the site of the crosslink. After ligation of adapter sequences to these ends, the sequences are determined by massively parallel sequencing. These adapter-ligation methods yield a rough approximation of crosslink location with best-case resolution of plus-or-minus ten nucleotides^9,21^, with the calculations of overall abundance biased by the complex multi-step ligation and library preparation steps required prior to sequencing^8,22^. In addition, commonly used crosslinking reagents and UV irradiation both have strong sequence and structural selectivity, such that observed crosslinks detect only a small fraction of intermolecular RNA interactions.

We recently introduced a strategy we call SHAPE-JuMP (for <u>s</u>elective 2’-<u>h</u>ydroxyl <u>a</u>cylation analyzed by <u>p</u>rimer <u>e</u>xtension and <u>ju</u>xtaposed <u>m</u>erged <u>p</u>airs)^23^ in which nucleotides in close three-dimensional proximity are crosslinked with a bi-reactive reagent (Fig. 1A, *left*). Initial experiments used the crosslinker *trans* bis-isatoic anhydride (TBIA) (Fig. 1B, *left*). TBIA is a SHAPE reagent and, as such, reacts with the 2’-hydroxyl group of unconstrained nucleotides, largely independent of nucleotide identity^24^. In SHAPE-JuMP, sites of crosslinking are recorded in a *single* direct step using an engineered reverse transcriptase (RT)^25^ that “jumps” across the crosslink during reverse transcription, creating a deletion in the resulting cDNA^23^. Deletion sites, and thus the positions of crosslinked nucleotides, are identified by massively parallel sequencing and alignment of the deletion-containing sequences. To control for non-crosslink-mediated deletions, an experiment is performed in parallel with a reagent that yields mono-adduct containing RNAs (Fig. 1A, *right*). This control experiment is performed with isatoic anhydride (IA), a molecule with a structure similar to TBIA, but with only one reactive moiety (Fig. 1B, *right*). The JuMP strategy provides, in principle, a very simple, direct and experimentally concise readout of sites of crosslinking in RNA. Nonetheless, as currently implemented, there are important limitations: The crosslink-jumping RT enzyme generates cDNAs with high levels of internal mutations, complicating accurate alignment; the “landing” site may be several nucleotides away from the site of the crosslink; and crosslinks are not always jumped consistently. We therefore developed a bioinformatic pipeline, ShapeJumper, to process SHAPE-JuMP sequencing data with the goal of mitigating these limitations.

**Figure 1:**
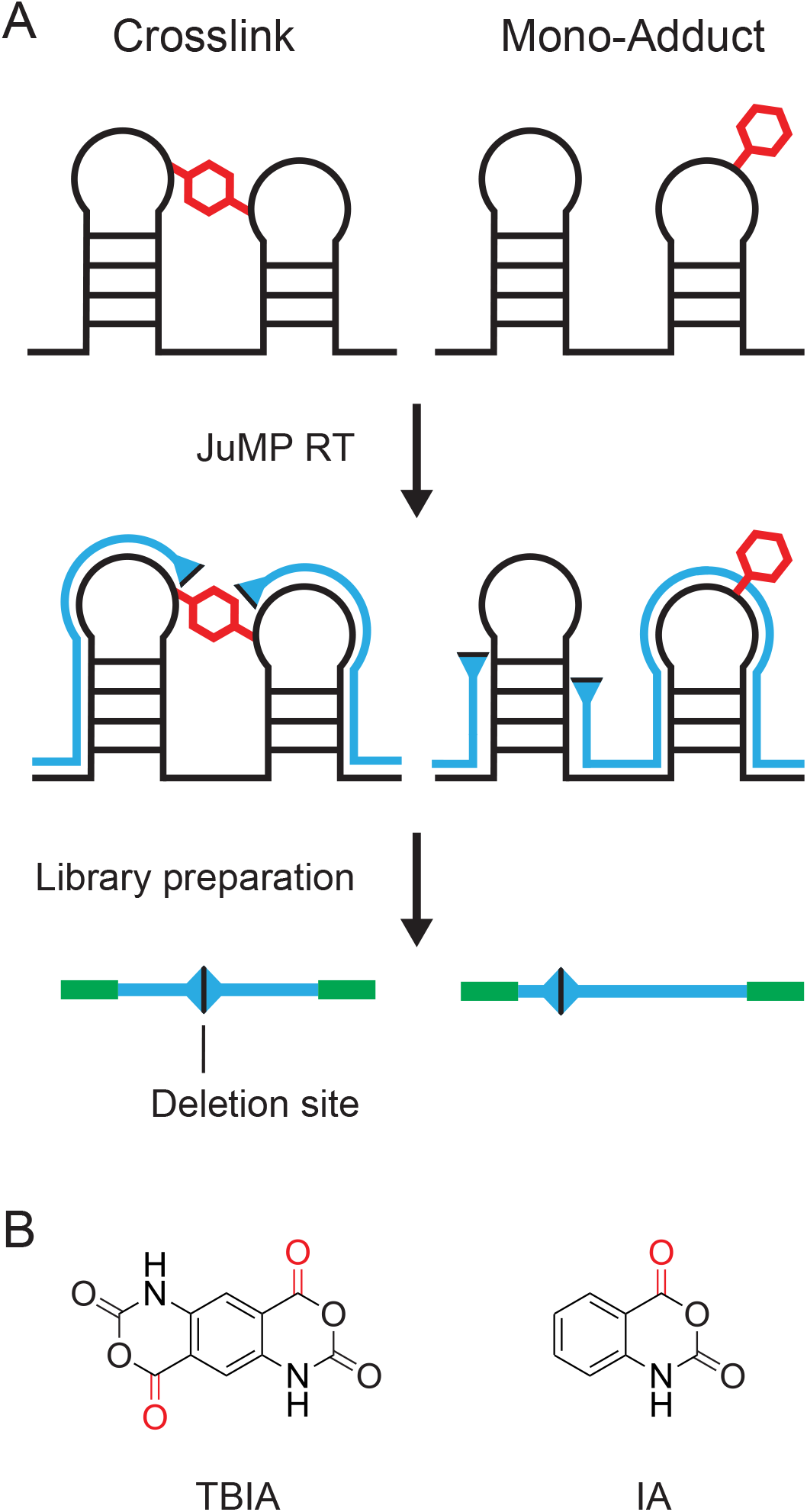
SHAPE-JuMP experimental overview. (**A**) RNA is crosslinked with a bi-functional SHAPE or other reagent, and the site of crosslinking is recorded as a deletion in the cDNA generated by reverse transcription under specialized RT conditions (*left*). In parallel, a control reaction that induces a mono-adduct (or no adduct) in the RNA is used to provide a control for non-crosslink-induced deletions (*right*). The cDNA is sequenced to identify deletion sites. (**B**) Examples of SHAPE reagents that form RNA crosslinks (TBIA, *left*) and monoadducts (IA, *right*). TBIA-dependent crosslinks, more frequent than the IA background, report proximal through-space interactions in RNA.

The ShapeJumper pipeline identifies crosslinked nucleotides from sequencing data (Fig. 2). Sequencing reads are first processed to remove low per-nucleotide quality scores and to merge overlapping reads. Reads are aligned^26^ with optimized parameters, as described in this work. The resulting alignment file is analyzed with a custom algorithm to identify deletion sites; during this process ambiguous deletions are removed and exact alignments are enforced at deletion sites to improve accuracy. Deletion rates are then normalized by read depth, and background rates for a non-crosslinked control are subtracted to correct for crosslink-independent deletions. ShapeJumper works well for most classes of crosslinking strategies, including SHAPE-based methods (TBIA), psoralen reagents, and UV irradiation.

**Figure 2:**
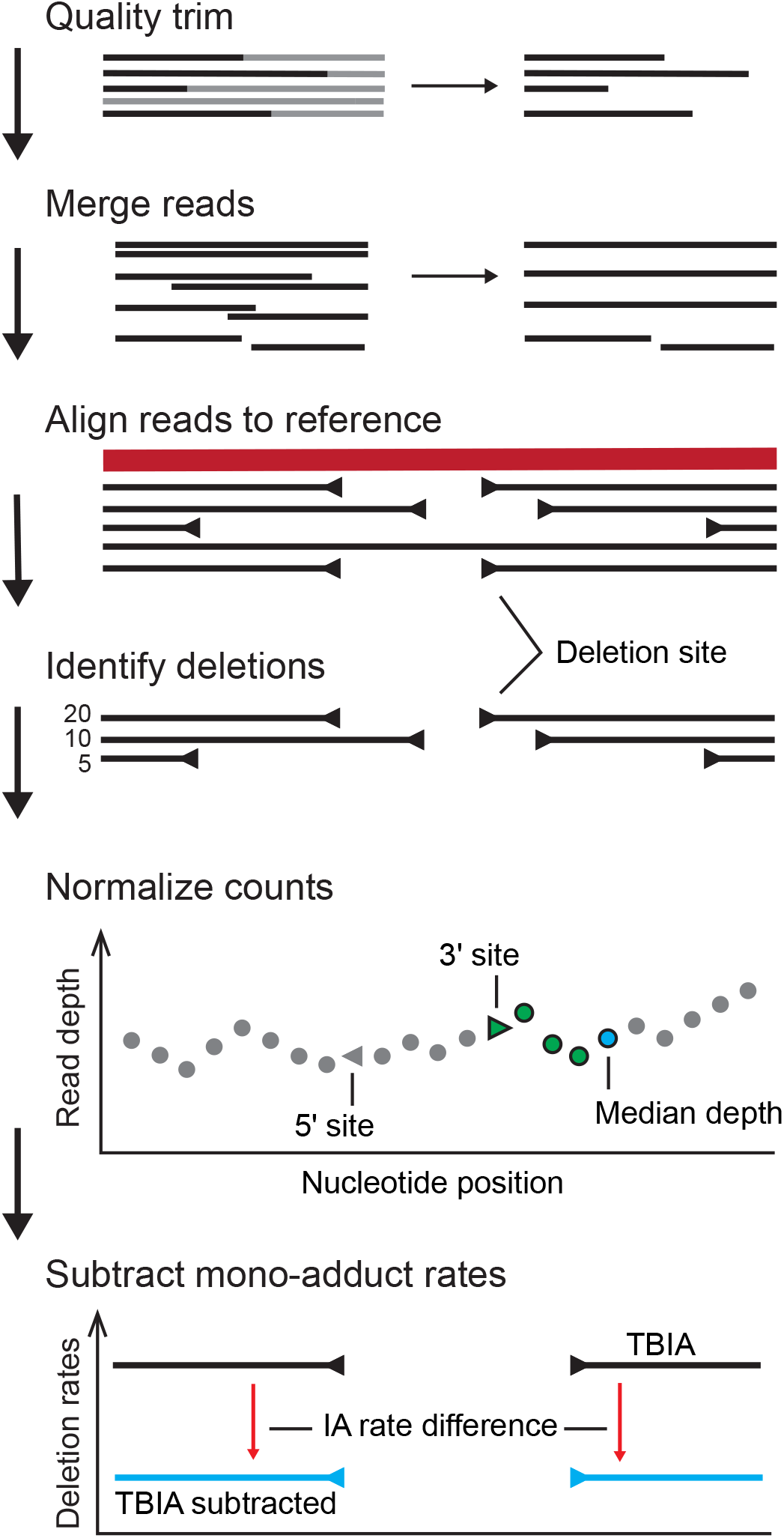
ShapeJumper overview. SHAPE-JuMP sequencing reads are processed for read quality, and paired reads (if used) are merged. Reads are aligned to a reference sequence, creating an initial set of candidate deletion sites. Candidate deletion sites are either identified from an alignment directly or inferred from two alignments separated by unaligned reference sequence. Deletion rates are normalized by the median read depth over the 5 nucleotides downstream of the 3’ deletion site. Normalized rates are obtained by subtracting mono- or noadduct rates.

## Results

### A preferred aligner for deletion analysis

Aligning SHAPE-JuMP derived reads accurately is a unique problem. Individual reads may or may not have a deletion, the deletions may vary in length, and the rates of occurrence of deletions vary. The RT enzyme currently used in the SHAPE-JuMP strategy has the special ability to read across crosslinked sites but also has a high non-crosslink-related per-nucleotide mutation rate of 3-4%^23^, which makes alignment challenging. No aligner has been specifically designed or optimized to operate with this type of complex data. We evaluated five aligners for use in the SHAPE-JuMP pipeline. BLAST, a sequence-comparison-focused algorithm, was selected as an example of a basic hash-table-based aligner^27,28^. YAHA, also hash-based, was selected because it was optimized to detect genomic structural variants, including deletions^29^. Hash-table aligners are slow but perform exhaustive searches of sequence space^30,31^. We also evaluated three aligner programs based on suffix/prefix tries (based on the Burrows Wheel Transform algorithm). These aligners are faster and thus better equipped to process large numbers of inputted reads^30^. Bowtie 2 was evaluated for its ability to process gapped alignments and accept mismatches^32^. BWA-MEM^26^ also allows for gapped alignments, is designed to handle sequencing errors robustly, and is optimized for reads of 100 to 1000 nucleotides. STAR was examined because it is an effective splice-site detection aligner^31,33^, which share some similarity with SHAPE-JuMP deletions. Aligner programs were assessed using their default parameters, except for small changes to Bowtie 2 and STAR (see Methods).

We evaluated the ability of these aligners to detect SHAPE-JuMP deletions using datasets of synthetic sequencing reads designed to mimic SHAPE-JuMP sequencing reads that contained known deletion locations. These datasets were designed specifically to contain sequences that mirrored those observed in experimental SHAPE-JuMP reads, performed with the RT-C8 enzyme^23^. Two synthetic read datasets were created, a deletion set and a deletion-insertion set. Both datasets consist of reads with randomly placed deletions. The deletion-insertion set contained deletions with additional random insertions of 1 to 9 nucleotides within the deletion. The frequency of each insertion length was sampled from a set of experimental reads. Mutations included mismatches, single-nucleotide insertions, and single-nucleotide deletions, each at levels proportional to their occurrence in experimental reads. These synthetic reads were analyzed using each of the five aligners, and the accuracies of the alignments were assessed by binning the observed deletions into one of three categories (Fig. 3A): An exact match defined as an alignment that predicts the site of the deletion correctly; close matches for which predicted 5’ and 3’ borders of the deletion are within three nucleotides of the actual site; and incorrect alignments that exceeded these limits. BLAST had the highest level of exact and close matches, but also had the highest level of incorrectly predicted deletion sites (Fig. 3B). STAR also had a high level of exact and close matches for the deletion read set but very few deletions were accurately predicted in the deletion-insertion set. Overall, BWA-MEM was the best performer in this analysis for accurately identifying sites of deletions without introducing a bias against detecting deletions in sequences containing deletion-insertions. BWA-MEM was thus used as the aligner in the SHAPE-JuMP pipeline.

**Figure 3:**
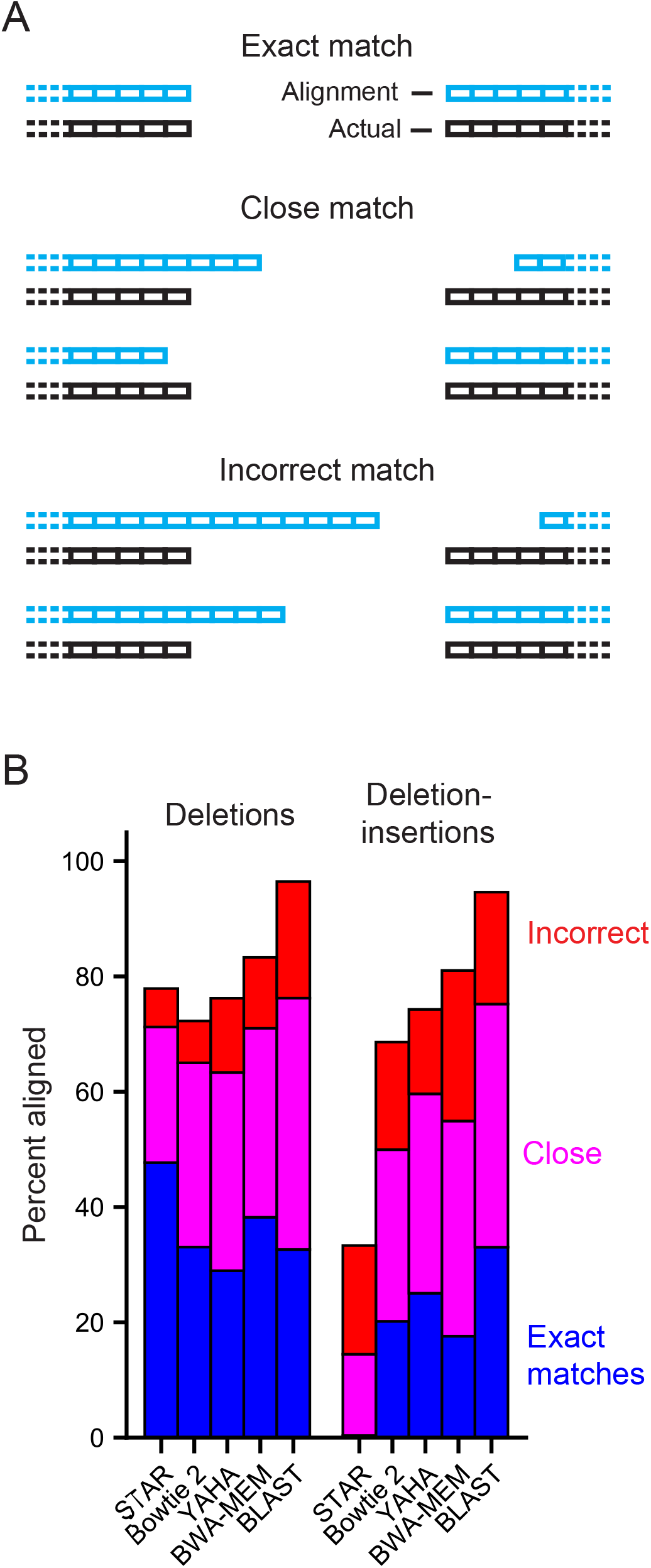
Accuracy analysis for candidate aligners. (**A**) Categories of aligned deletions. (**B**) Analysis of performance of a representative set of aligners on synthetic read datasets. Alignments were performed using two sets of synthetic data: containing deletions and deletioninsertions. The deletions set consists of reads with a randomly placed deletion whereas the deletion-insertions dataset also contained a 1-9 nucleotide sequence insertion at the deletion site. Both synthetic datasets contain point mutations reflective of those observed in experimental reads. Each synthetic dataset contained one million reads generated from an RNase P reference sequence. Match categories reported as percentage of total reads in each category.

### Alignment and deletion detection optimization

BWA-MEM was incorporated into a proto-ShapeJumper pipeline and was optimized to address the low positive-predictive value (ppv) for a substantial subset of deletions in the synthetic deletion dataset (Fig. 4A). Here ppv is defined as the fraction of predicted deletions that occur in the synthetic data set, at a given set of coordinates. Default BWA-MEM scoring parameters^26^ were altered as follows: (*i*) the score penalty for mismatches (–B) was lowered from 4 to 2 to account for the high mutation rate of reads; (*ii*) the deletion score penalty (–O) was decreased from 6 to 2 to accommodate high mutation rates and to promote alignment of longer deletions; and (*iii*) the scoring threshold (–T) was lowered from 30 to 15 and the initial seed (–k) shortened from 19 to 10 to allow reads with short sequences flanking a deletion to be aligned. These changes substantially increased deletion site calling accuracy, increased the number of deletions in short sequencing reads that could be aligned, and reduced the fraction of deletions that were incorrectly aligned.

**Figure 4:**
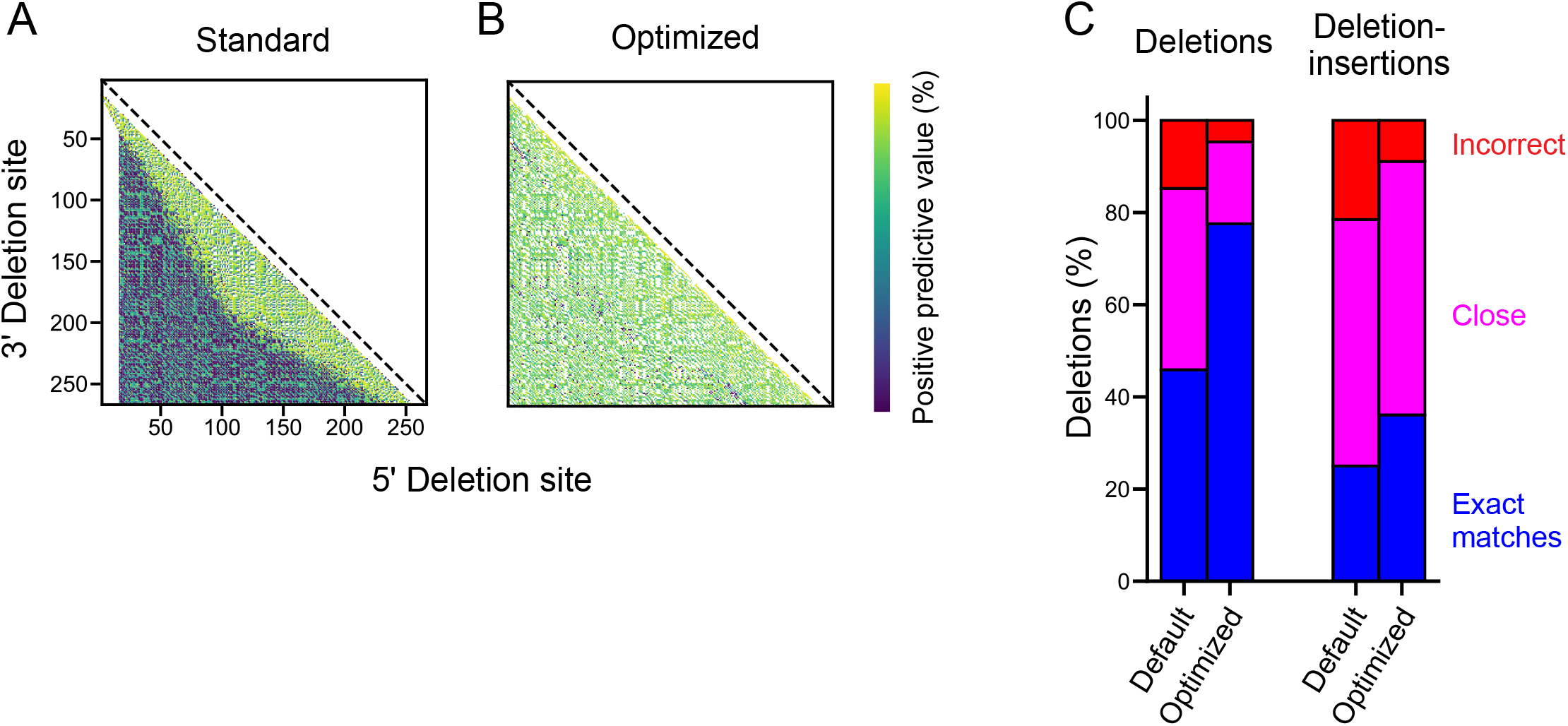
Alignment optimization. Interaction maps for deletion sites identified from the deletion dataset of synthetic reads for BWA-MEM alignment with (**A**) default parameters, (**B**) optimized algorithm. The optimized analysis incorporates custom BWA-MEM parameters, ambiguous site removal, and exact edge matching. Points correspond to specific 5’ and 3’ deletion sites and are colored by the percent of total deletion sites correctly mapped to a specific nucleotide pair (see scale). (**C**) Summary of accuracies pre- and post-optimization for synthetic deletion (*left*) and deletion-insertion (*right*) datasets.

Following these scoring alterations, there remained a systemic bias in the alignment of ambiguous deletions, defined as deletions where one site cannot be uniquely identified because the same nucleotide is present at both sides of the deletion (Fig. S1A). The scoring function used during alignment extension from the initial seed leads to the ambiguous nucleotide always being aligned before the gap opening, resulting in a directional bias in deletion-site detection.

ShapeJumper therefore removes ambiguous deletions, which results in more accurate alignment of the neighboring, unambiguous deletions (Fig. S1B), and results in a roughly 20% increase in exact match detection.

Deletion-insertions also exacerbate inaccurate deletion site assignments, if the insertion includes nucleotides matching the reference sequence within the region of a deletion (Fig. S1C). To mitigate insertion-induced misalignments, edge matching was enforced for all deletion sites such that three nucleotides on both sides of the deletion site are required to exactly match the reference. If this is not the case, the deletion site is moved one nucleotide to the exterior, and the removed nucleotide is identified as an insertion in the alignment. This process is repeated until all three nucleotides at the deletion site match the reference. Enforcing exact edge matching notably increased the accuracy of short deletion detection without compromising overall deletion detection (Fig. S1D). The combined effect of these optimizations, custom BWA-MEM parameters, ambiguous deletion removal and exact edge matching, substantially increases deletion site detection accuracy (Fig. 4B, 4C).

### Experimental data-driven optimization of pipeline

After optimizing the pipeline with synthetic data, the proto-ShapeJumper pipeline was used to process experimentally generated SHAPE-JuMP reads obtained from analyses of a set of small to large RNAs (158-412 nts): the P546 group II intron domain, M-Box riboswitch, Varkud satellite ribozyme, RNase P catalytic domain, and group II intron^23^. Quality filtered and merged reads were aligned, the resulting alignments parsed to identify deletion sites, and deletion rates were normalized by the median read depth of the 5 nucleotides downstream of the 3’ deletion site (Fig. 2). Normalization also enables comparison between samples, including the non-crosslinked (IA) control. The normalized deletion rates observed in the non-crosslinked experiment are subtracted from those observed in the crosslinking experiment to control for non-crosslink-induced deletions. Normalization thereby also removes outliers with high deletion rates (Fig. S2). After this background subtraction step, the most frequent deletions more accurately reflect a holistic view of proximal RNA-RNA interactions (Fig. S2C). Background normalization also yields increased area under curve (AUC) in receiver operating characteristic (ROC) curves for through-space interactions within 15 Å of each other for an RNA with complex higher-order structures (Fig. S2D).

Long insertions in insertion-deletions are prevalent in experimental data and can contribute to alignment error. For example, for the RNase P RNA, approximately 50% of deletions contain an insertion of at least one nucleotide (Fig. *S3blue*). Insertions were a substantial source of error in the synthetic read alignments, as evidenced by the difference in accuracy for predicting deletions compared to deletion-insertions (Fig. 4C). Insertion length and deletion site assignment error are correlated. (Fig. S3, *red*). ShapeJumper therefore removes reads containing a deletion with an insertion size greater than 10, decrementing the count of deletions found at that site. Insertions longer than 10 nucleotides are infrequent so their removal had a small effect on the total number of deletions detected (Fig. S3, *blue*), and moderately improved deletion site detection.

Finally, experimental SHAPE-JuMP data were analyzed to identify additional features that might improve the precision of detecting proximal interactions. The RT enzyme jumps the crosslink in the 3’ to 5’ direction (Fig. 5A), and it is possible that the nucleotides that physically form crosslinks are downstream of the 5’ site or upstream of the 3’ site. We examined this possibility by shifting the assigned 5’ and 3’ sites 0 to 5 nucleotides downstream and upstream, respectively, and examined the effect of these shifts on known through-space inter-nucleotide distances. Shifting the 5’ crosslink site 2-nucleotides upstream both increased the detection rate for tertiary interactions and decreased the through-space distance of reported interactions (Fig. 5C, S4).

**Figure 5:**
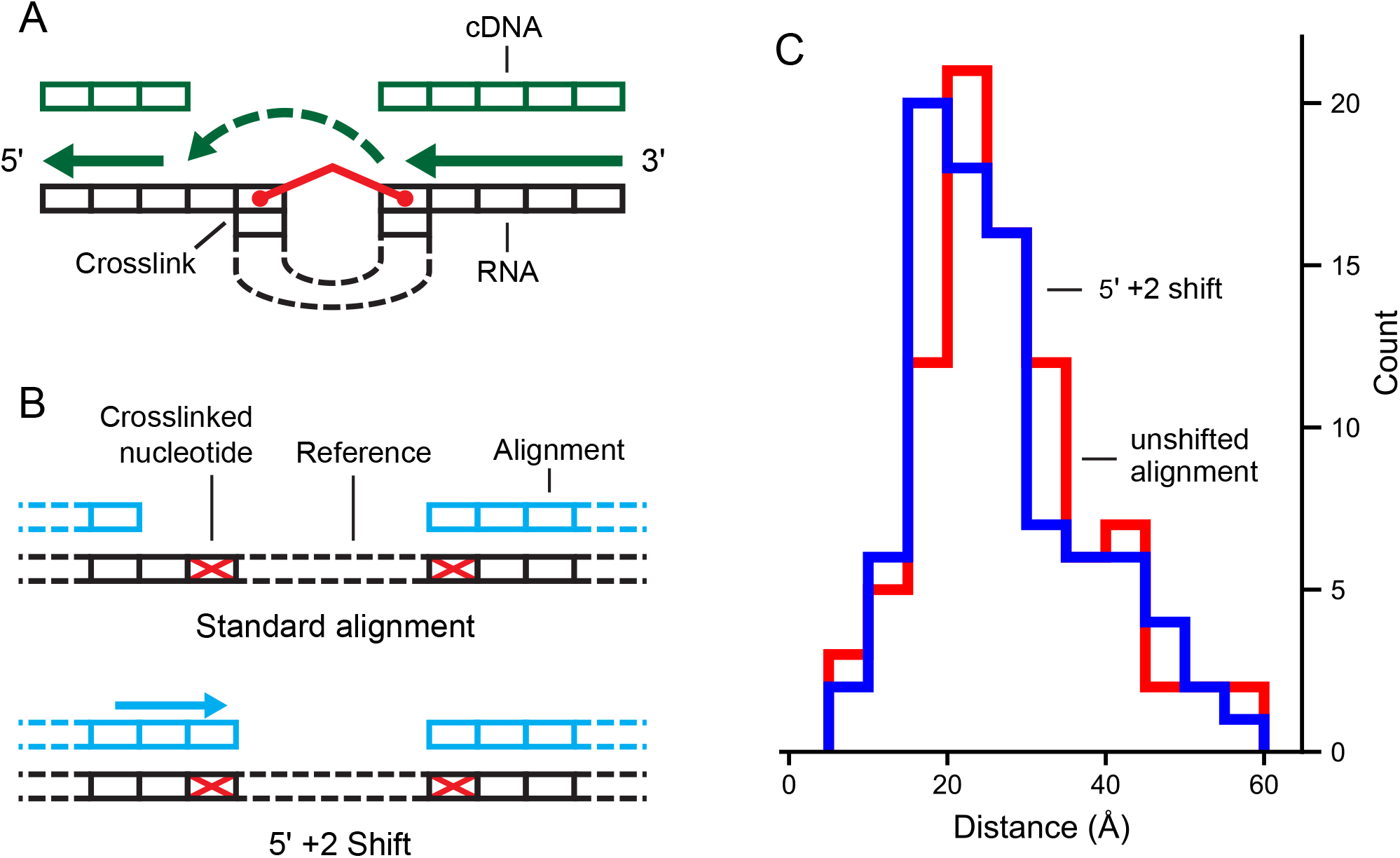
Effect of shifting deletion site assignment on through-space distance. (**A**) Relationship between crosslinked RNA and RT jumping. Directionality of reverse transcription and crosslink-induced steric hindrance can yield an offset at the 5’ deletion site relative to crosslink position. (**B**) Deletion site adjustment to compensate for the mechanism of RT jumping. (**C**) Distance distribution of RNase P RNA SHAPE-JuMP data for unshifted (red) and shifted (blue) deletion assignments. Through-space distances are shown for the deletions corresponding to the most frequent 3% of deletions. Datasets were processed using the fully optimized ShapeJumper pipeline with the exception of the shift or not for the RT landing mechanism.

### Applications of ShapeJumper

ShapeJumper includes useful tools for troubleshooting and visualizing the results of RNA crosslinking experiments. ShapeJumper tools report the distribution of deletion rates and filter deletions by sequence length, which requires only the primary sequence of the target RNA. ShapeJumper calculates contact distances of deletions, defined as the distance between nucleotides after omitting nested helices, which provides a measure of proximity in secondary structure versus primary sequence space^34^. ShapeJumper also provides visualization tools that facilitate efficient assessment of the quality of a crosslink strategy or experiment. Deletions can be plotted, at any level of frequency, on a secondary structure diagram (Fig. 6A). Given a known or modeled three-dimensional structure, deletions can be visualized and colored by through-space distance (Fig. 6B). Three-dimensional distances can be plotted for a given deletion rate and compared to the distance distribution expected by chance (Fig. 6C).

**Figure 6:**
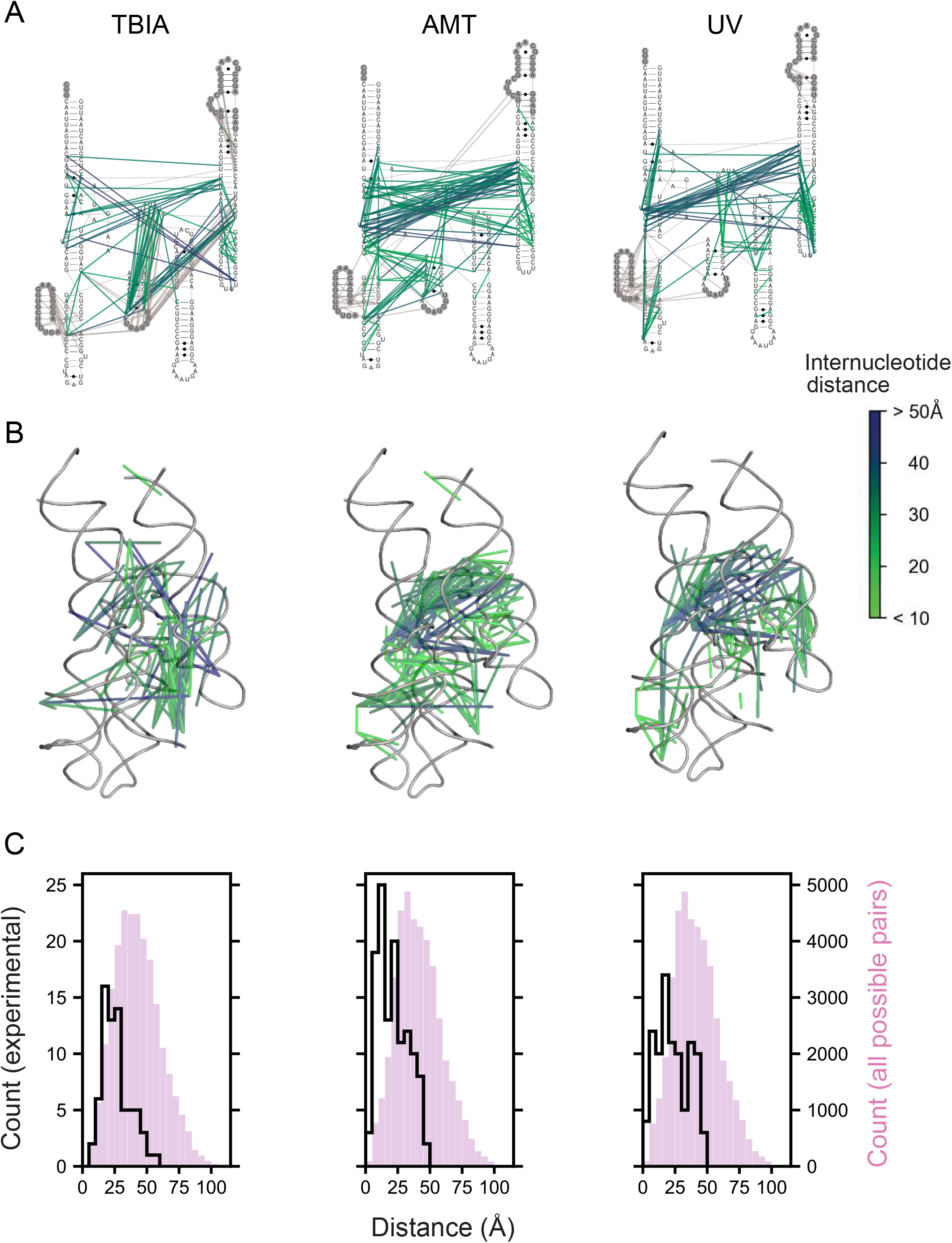
ShapeJumper measures deletions obtained from diverse crosslinkers. Columns illustrate ShapeJumper analysis of experiments performed with the SHAPE reagent TBIA^23^(*left*), the psoralen derivative AMT (*middle*), and short wavelength UV (*right*). Crosslinks were obtained with the RNase P catalytic domain RNA^43^; the 3% most frequent deletions are shown. (**A**) Deletions superimposed onto the secondary structure. Deletions observed in regions not visualized in the reference three-dimensional structure are gray. (**B**) Deletions superimposed onto a tertiary structure model. In panels (A) and (B), deletions are shown as lines and are colored by through-space distance between nucleotides. (**C**) Distance distribution of deletions. Distances as measured by crosslink-induced deletions are shown as lines; all possible distances are shown with magenta histograms. Distances were measured between ribose 2’-hydroxyl groups (*left*) or between central point of the nucleobase (*middle and right*).

The SHAPE-JuMP strategy and ShapeJumper software work for a wide variety of crosslinking reagents. We have successfully implemented ShapeJumper to evaluate RNA crosslinking experiments performed with TBIA, the psoralen derivative 4’-aminomethyltrioxsalen hydrochloride (AMT), and short wavelength UV (Fig. 6). The patterns of observed deletions vary, reflecting the distinct underlying chemistry of each reagent but, overall, clearly map proximal sites in the large RNase P RNA. We anticipate that most sequencing-based proximal-interaction identification methods^13–20^ can be processed and analyzed via ShapeJumper, yielding excellent performance in the accuracy of deletion assignment sites and rates.

## Discussion

### Deletion site identification optimization

In principle, crosslinking represents a simple and direct way to map through-space interactions in RNA. In practice, the power of RNA crosslinking has been difficult to realize because of numerous challenges in detecting sites of crosslinks accurately, and at nucleotide resolution. Identification of an RT enzyme that has the distinctive activity of extending cDNA synthesis through the sites of crosslinks in RNA, revealing these sites as deletions in the cDNA, is an important experimental advance. The cDNA signals are currently complex, however, as the RT enzyme yields cDNAs with internal mutations, the landing sites may be several nucleotides away from the site of the crosslink, and the crosslink may cause termination of polymerization. The ShapeJumper pipeline was designed to be aware of these challenges, to identify and quantify crosslink-mediated deletions, and to distinguish crosslink-induced deletions from other polymerase-mediated mutations.

Deletion rates in a SHAPE-JuMP experiment can vary substantially between RNA targets and it is therefore important to identify infrequent deletions. ShapeJumper attempts to maximally predict deletion sites by allowing low alignment-score thresholds (Fig. 4B). Deletion rate variation and polymerase-mediated sequence deletions complicate reproducibility. ShapeJumper addresses these complicating features by normalizing deletion rates by read depth (Fig. 2).

Finally, the deletion rates of a mono-adduct experiment are subtracted from crosslinked deletion rates to control for crosslink-independent RT-mediated deletions (Figs. 2 and S2C).

ShapeJumper was optimized to maximize detection accuracy for the 5’ and 3’ deletion sites. The crosslinker used for SHAPE-JuMP in our exploratory studies spans ~7 Å between active sites (Fig. 1B, *left column*). Crosslinked nucleotides should be similarly close in three-dimensional space. Misidentifying the deletion site by just one nucleotide increases the inferred distance by 10-15 Å^35^. We do observe a fraction, 4%, of distances of 45 Å or greater, which likely reflect a combination of false positive measurements, conformational dynamics in these large RNAs, or other features not reflective of internucleotide distances. Accuracies of five aligners were examined, using synthetic datasets with reads containing single-nucleotide mutations, deletions, and insertions, and insertions in the context of deletions (Fig. 3). Removing ambiguous deletions and forcing exact edge matching increased assignment accuracies (Fig. 4, S1). The net effect of our aligner choice and these optimization steps is a pipeline that accurately identifies sites of crosslinking, and thus RNA through-space interactions, as shown by analysis of data from a representative set of small and medium sized RNAs (Fig. S5) and for multiple classes of crosslinking experiments (Fig. 6). We note that parameter choices and optimization steps were tailored to the specific mutation and deletion characteristics of the RT-C8^25^ enzyme, characterized for SHAPE-JuMP^23^. We think the algorithm developed here will be effective for alternative jumping polymerases and reagents identified in the future, with only minor modification or optimization of the ShapeJumper pipeline.

### Perspective

Melding either per-nucleotide RNA chemical probing or through-space crosslinking experiments with a readout by massively parallel sequencing enables analysis of RNA structure with unprecedented throughput and impressive detail. Among many useful applications, SHAPE-JuMP can be used to map through-space interactions in large, complex RNAs (Fig. 6), and identify restraints useful for three-dimensional RNA structure modeling^23^. However, it is a challenge to convert the direct results of chemical probing or crosslinking into a form readable by massively parallel sequencing. The ongoing transition from experimentally complex -seq class experiments to much more direct mutational profiling (MaP) has simplified the experiment and increased the accuracy of per-nucleotide chemical probing^3,7,36^. Analogously, a transition from complex adapter-ligation protocols to direct JuMP experimental readouts appears poised to transform experiments that measure through-space RNA-RNA interactions via crosslinking. A key to both MaP and JuMP readouts is software that carefully accounts for the idiosyncrasies of these experiments. ShapeJumper detects deletions resulting from crosslink jumping – from which RNA-RNA interactions can be inferred – with near-nucleotide resolution. The pipeline is easy to implement, requires little to no user input after execution, and works for diverse crosslinking reagents. SHAPE-JuMP and ShapeJumper inaugurate new platforms for efficient detection and analysis of through-space interactions for diverse RNA targets.

## Methods

### Jumping RT enzyme

Data analyzed in this work were generated by the RT-C8 enzyme, developed by directed evolution using a compartmentalized bead labelling strategy^25^.

### ShapeJumper pipeline

ShapeJumper is a Bash script that executes multiple python programs and is executable on most UNIX platforms. Inputs are Illumina sequencing reads of crosslinked and non-crosslinked samples in FASTQ format and a reference sequence file in FASTA format. By default, a text file with deletion locations and normalized, background-subtracted rates is output. Ambiguous deletions and deletions with insertions of 10 nucleotides or greater are removed, exact edge matching of deletion sites is enforced, and the final reported deletions have undergone a 5’ +2 shift. Alignment and processing parameters can be varied, as described in the included documentation. Additional python tools are provided for analysis of measured deletions in terms of their distribution at the levels of sequence and secondary and tertiary structure. Python 2.7 and necessary third-party packages are available from the Conda package manager (https://conda.io/docs/). The following algorithms are used in the pipeline: ShapeMapper v2^37,38^ is used to trim reads by base-call quality; FLASH^39^ is used for merging overlapping reads; BWA-MEM^26^ is used to align reads to the reference sequence; and PYMOL (https://pymol.org/) and Biopython^40,41^ are used for tertiary structure analysis.

Raw sequencing reads are trimmed by base-call quality using the read trimmer program, ShapeMapper read trimmer, part of ShapeMapper v2^38,42^. Quality scores for each nucleotide in a read are scanned from 5’ to 3’. When the first set of 5 nucleotides with an average quality score below 20 is identified, it and all downstream nucleotides are removed from the read. Reads shorter than 25 nucleotides, post trimming, are removed. The resulting trimmed reads are then merged with their pair mate using FLASH^39^, which increases quality scores in the overlapping region. Reads that do not overlap are not removed. The quality trimmed and pair mate merged reads are aligned using BWA-MEM^26^ with the parameters optimized in this work: Gap open penalty (–O) of 2, mismatch penalty (–B) of 2, minimum seed length (–k) of 10, score threshold for output alignments (–T) (lowered to) 15. All reads were parsed from cigar strings. Merged and unmergeable reads are parsed and aligned separately, their outputs combined, and duplicate deletions are removed.

Short deletions are directly identified by the aligner. Longer deletions generally result in two alignments per read, one each for the sequence upstream and downstream of the deletion; deletions are identified as the intervening reference sequence between the two alignments that did not align to the read. Multiple deletions can be detected in a single read. Deletions shorter than 10 nucleotides or deletions with an insertion of greater than 10 nucleotides are ignored. The 3 nucleotides upstream and the 3 nucleotides downstream of the deletion site are required to exactly match the reference. If there is a mismatch, the deletion site is shifted until there is an exact match. If these shifts involve more than 10 nucleotides total, the deletion is not reported.

Deletion counts are normalized by the median read depth of the 5 nucleotides immediately downstream of the 3’ deletion site. The normalized rates of the mono-adduct control sample are subtracted from the normalized deletion rates of the crosslinked sample. Deletions detected only in the crosslinked sample are retained. Finally, 5’ deletion sites are shifted two nucleotides in the 3’ direction (Fig. 5B). The final deletion data set is reported as each deletion 5’ and 3’ site, with the normalized and subtracted rate of occurrence.

### Aligner evaluation

We emphasize from the outset that current aligners were not designed for our application; nonetheless, most tested aligners could be used to interpret JuMP data in a useful way. In general, we used each aligner with default or near-default parameters, and improvements to the non-selected aligners is likely possible with additional parameter changes. Aligners were tested using two datasets, each comprised of 1,000,000 computationally generated synthetic reads: a deletion set and a deletion-insertion set. Both synthetic read sets were generated by placing deletions randomly in the sequence for the RNase P catalytic domain^43^; the sequence included flanking structure casettes^24^ but deletions were not placed in the structure cassette sequences. The deletion-insertion set contains deletions generated in this manner, but the deletions also contained an additional insertion. The insertion lengths were randomly sampled from the distribution of insertion lengths observed from a SHAPE-JuMP experiment using the RNase P RNA^23^ (Fig. S3, *blue line*). Reads in both sets were randomly mutated at single nucleotides at an overall rate of 3.75%; of these mutations, 3% were insertions, 26% were deletions, and 71% were single-nucleotide changes. These rates and ratios mimic the activity of the jumping RT used in this work.

Reads were aligned using the default parameters for each tested aligner with two exceptions. For Bowtie 2, the alignments reported parameter (-k) was set to 2 to enable detection of longer deletions. For STAR, the minimum intron size was set to 10 and the non-canonical junction penalty was lowered to −4 to increase the rate at which deletions were identified at exon junctions; this change was explored to take advantage of splice-site reporting in STAR and to possibly forgo the need to parse deletion sites from SAM files.

The resulting alignments were parsed for deletion-site locations. Locations were then compared to the known deletion sites encoded in the synthetic reads. Each alignment was binned into one of three deletion identification categories: exact matches, where the aligned deletion sites exactly match the encoded deletion sites; close matches, where both of the aligned deletion sites are within 3 nucleotides of the encoded site; and incorrect matches, where one or both of the aligned deletion sites are more than 3 nucleotides from the encoded site. The same synthetic reads and matching criteria were used to evaluate and develop custom BWA-MEM parameters.

As part of the BWA-MEM optimization strategy, a third increasing-insertion-length synthetic read dataset was created to evaluate the effect of insertion length on deletion-site detection accuracy. This dataset consisted of deletions that contain insertions of lengths ranging from 0 to 30. 100,000 reads were synthesized for each insertion length. Reads were created from an RNase P catalytic domain reference sequence^24,43^ and mutated as described above. The increasing-insertion-length read dataset was aligned using BWA-MEM with ShapeJumper parameters. The resulting alignments were analyzed for deletion site accuracy at each insertion length (Fig. S4, *red*).

### Structure datasets

TBIA and IA SHAPE-JuMP datasets were collected previously^23^. The two reactive sites on TBIA react with half-lives of 30 and 180 sec; experiments are carried out for 15 min, equal to 5 half-lives of the slower reaction. Short-wavelength UV and 4’-aminomethyltrioxsalen hydrochloride (AMT) data sets were generated using a modified version of the SHAPE-JuMP protocol. Briefly, 15 pmol *in vitro* transcribed RNase P RNA was heat denatured for 1 minute and placed on ice. The RNA was incubated in folding buffer [100 mM HEPES (pH 8.0), 100 mM NaCl, 10 mM MgCl_2_] at 37 °C for 30 minutes, divided into three 18 μL aliquots, and transferred to amber tubes. One aliquot was treated with 1/9 volume 2 mg/mL AMT (Sigma-Aldrich A4330), dissolved in water, to yield a final concentration of 200 ng/mL AMT. The other two aliquots were treated with the same volume of water, one to serve as a control and the other to be crosslinked with short wavelength UV. The samples were incubated at 37 °C for an additional 15 minutes then placed on ice for crosslinking. The control and AMT samples were exposed to 365 nm (UVP CL1000; 10 cm from light source) for 30 minutes. The short-wavelength UV sample was exposed to 295 nm (UVP Handheld UV lamp, 6 W; 15 cm from light source) for 15 minutes. The RNA was purified by size-exclusion chromatography (G50 column, GE Healthcare) and kept on ice until reverse transcription. Reverse transcription was then performed using target-specific primers under SHAPE-JuMP conditions^23^ to produce a cDNA library. PCR was used to amplify cDNAs and to incorporate unique sequence barcodes^23^. Barcoded samples were sequenced (Illumina MiSeq instrument; 500 cycle v2 reagent kit). All datasets were analyzed with default ShapeJumper parameter sets (as developed in this work); for psoralen and UV crosslinking, analysis scripts were updated to define the center of the nucleobase as the site of crosslinking.

### Tertiary contact ROC curve analysis

All receiver operating characteristic (ROC) curve analyses used the same classifier, the set of nucleotide pairs with a three-dimensional distance less than 15 Å, and a contact distance greater than 10, where contact distance is defined as the sequence length between two nucleotides according to the secondary structure model when nested helices are skipped. This classifier was chosen as a way to approximate pairwise interactions that correspond to tertiary interactions. The true positive rate is defined as the fraction of ShapeJumper reported contacts with deletion rates above a given threshold that match this definition of tertiary contacts. The false positive rate is defined as the fraction of ShapeJumper contacts with deletion rates above a given threshold that do not correspond to a tertiary contact.

Deletion-site shifts were assessed using data from previously described SHAPE-JuMP experiments performed on five small RNAs^23^ with known three-dimensional structures: the *T. thermophila* group I intron P546 domain^44^, *B. subtilis* M-box riboswitch^45^, the *N. intermedia* Varkud satellite ribozyme^46^, the catalytic domain of *B. stearothermophilus* RNase P^43^, and the *O. iheyensis* group II intron^47^. To assess the effect of shifting the assignment for the 5’ and 3’ sites of crosslinking, SHAPE-JuMP reads were analyzed using default ShapeJumper parameters, and the resulting deletion junction sites were shifted by 0 to 5 nucleotides downstream of the 5’ deletion site and/or 0 to 5 nucleotides upstream of the 3’ deletion site. Shifted contacts were assessed using ROC curves and mean area under curve (AUC). ROC curve analysis was also carried out to assess the effect of each ShapeJumper analysis step (Fig. S5).

### Three-dimensional RNA structure modeling

One application of SHAPE-JuMP and the ShapeJumper pipeline is to identify restraints useful for three-dimensional structure modeling of complex RNAs. A detailed algorithm for SHAPE-JuMP based structure modeling is provided in a companion manuscript^23^.

## Supporting Information

Alignments and raw and processed deletions are available at https://weekslab.com/. The ShapeJumper pipeline is available at https://github.com/Weeks-UNC/ShapeJumper_V1. Structure analysis and visualization scripts available at https://github.com/Weeks-UNC/StructureAnalysis. Raw sequencing data from SHAPE-JuMP experiments^23^ can be obtained from sequence read archive <u>PRJNA687281</u>.

## Acknowledgements

We thank C.A. Lavender for the initial design of the deletion parsing script and M.J. Smola for initial design of the tertiary structure analysis scripts. This work was supported by the US National Institutes of Health (R35 GM122532 and R01 AI068462 to K.M.W. and R01GM101237 to A.L.). T.W.C. and C.A.G. were supported in part by NIH training grants, in Bioinformatics (T32 GM067553) and Molecular Biophysics (T32 GM008570), respectively.

---

**Figure S1:**
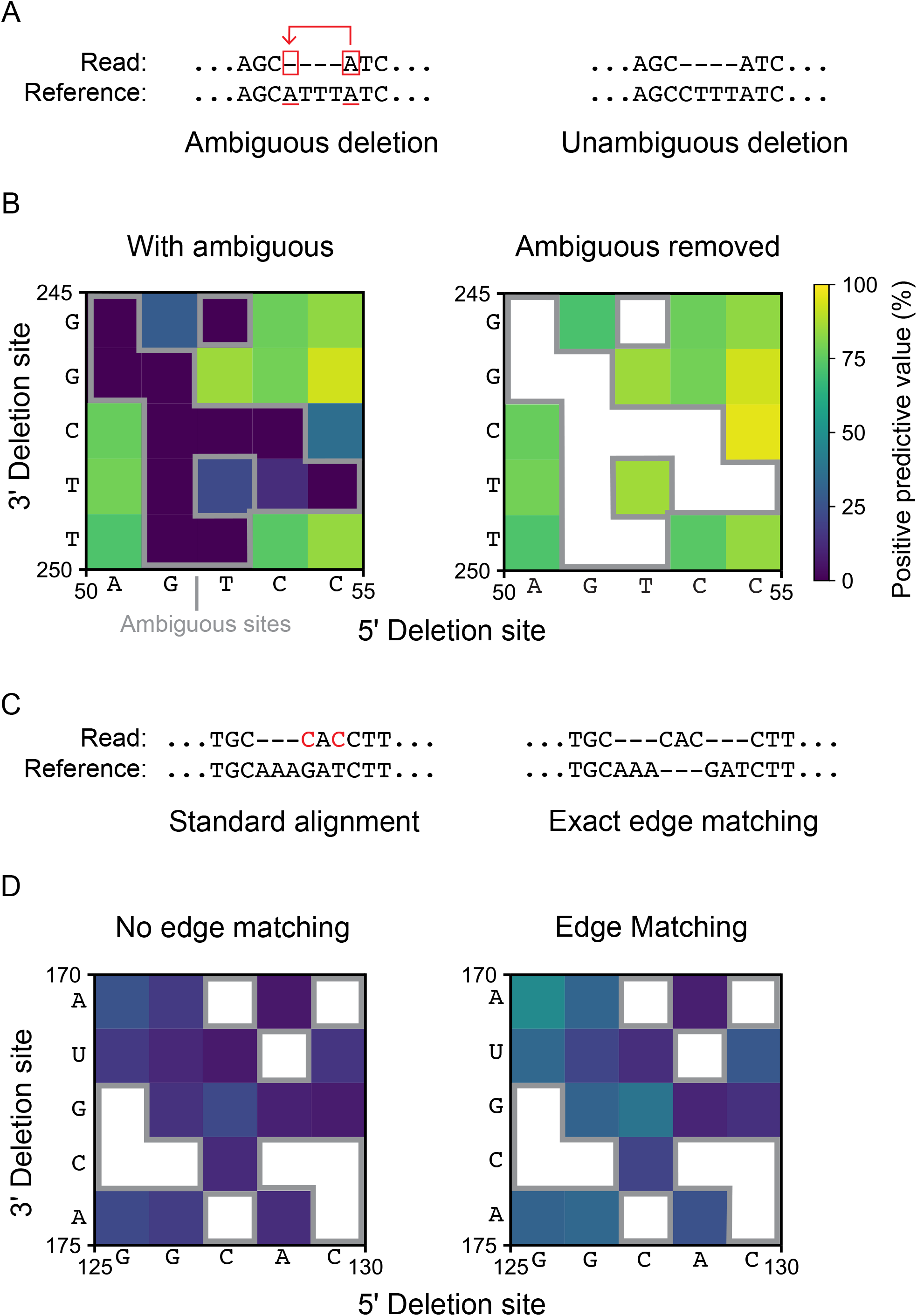
Effects of removing ambiguous deletions and enforcing exact edge matching. (**A**) Definition and example of an ambiguous deletion. An ambiguous deletion cannot be mapped to a unique site (*left*); an unambiguous deletion can (*right*). (**B**) Representative contact map of deletion sites from synthetic deletion read alignments containing (*left*) and without (*right*) ambiguous deletions. Ambiguous deletions enclosed in gray outline. Sites with no mapped deletions are white. Note extensive purple regions (0% ppv) are eliminated by removing ambiguous deletions. (**C**) Effect of enforcing exact edge matching (of 3 nucleotides) at a deletion site that also contain an insertion. (**D**) Representative contact map of deletion sites from synthetic deletion-insertion read alignments with ambiguous deletions removed without (*left*) and with (*right*) edge matching. Ambiguous deletions sites are removed in both cases. All contact maps (**B**, **D**) are colored on the same scale by the percent of deletions correctly mapped to a specific nucleotide pair.

**Figure S2:**
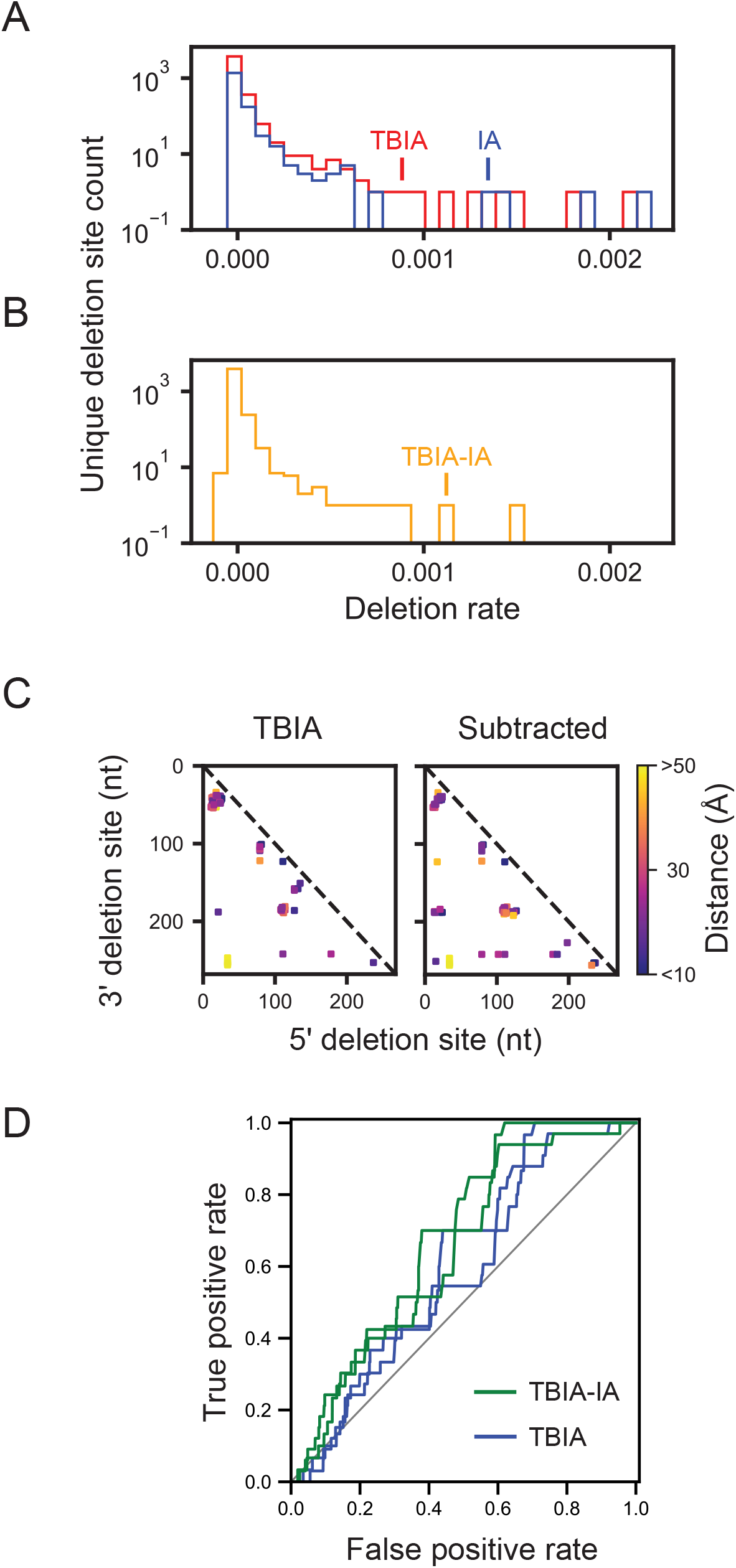
Improvement in TBIA-specific deletion rate measurement upon background subtraction. (**A**) Comparison of distributions of normalized deletion rates for crosslinked (TBIA) and mono-adduct (IA) RNase P RNA experiments. RNase P data used here show trends found in all RNAs examined to date. (**B**) Distribution of crosslink-induced deletion rates after mono-adduct subtraction. (**C**) Deletion sites corresponding to the 3% most frequent deletion rates, pre and post mono-adduct subtraction. Deletion sites are mapped onto the reference tertiary structure^43^ and colored by the three-dimensional distance separating the crosslinked nucleotides. (**D**) Ability of ShapeJumper to identify short distance interactions. ROC curve comparison based on TBIA-mediated crosslinking of the RNase P RNA^23^. Tertiary contact identification is shown for raw TBIA deletion rates (blue) and for TBIA deletion rates after subtraction by the IA control (green). Classifier: Inter-nucleotide distance less than 15 Å with a contact distance > 10 (see Methods).

**Figure S3:**
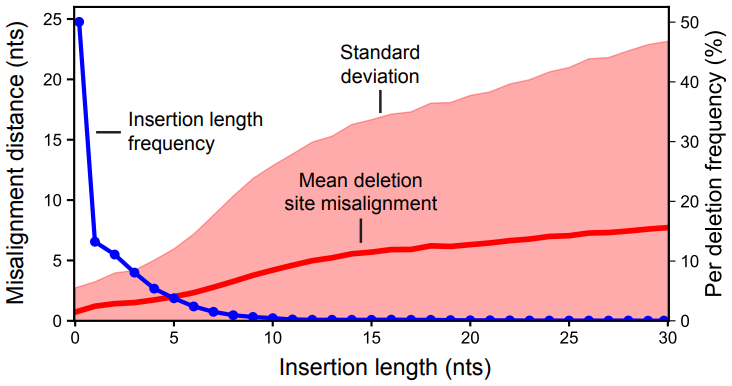
Insertion length effects on deletion site assignment accuracy and experimental deletion frequency. Misalignment distance is the sequence distance between assigned and known deletion end points. Mean misalignment distance (red line) as a function of insertion length in red. Standard deviation of misalignment is shown by red shading. Observed frequency of each insertion length in experimental SHAPE-JuMP RNase P data^23^ is shown with blue line.

**Figure S4:**
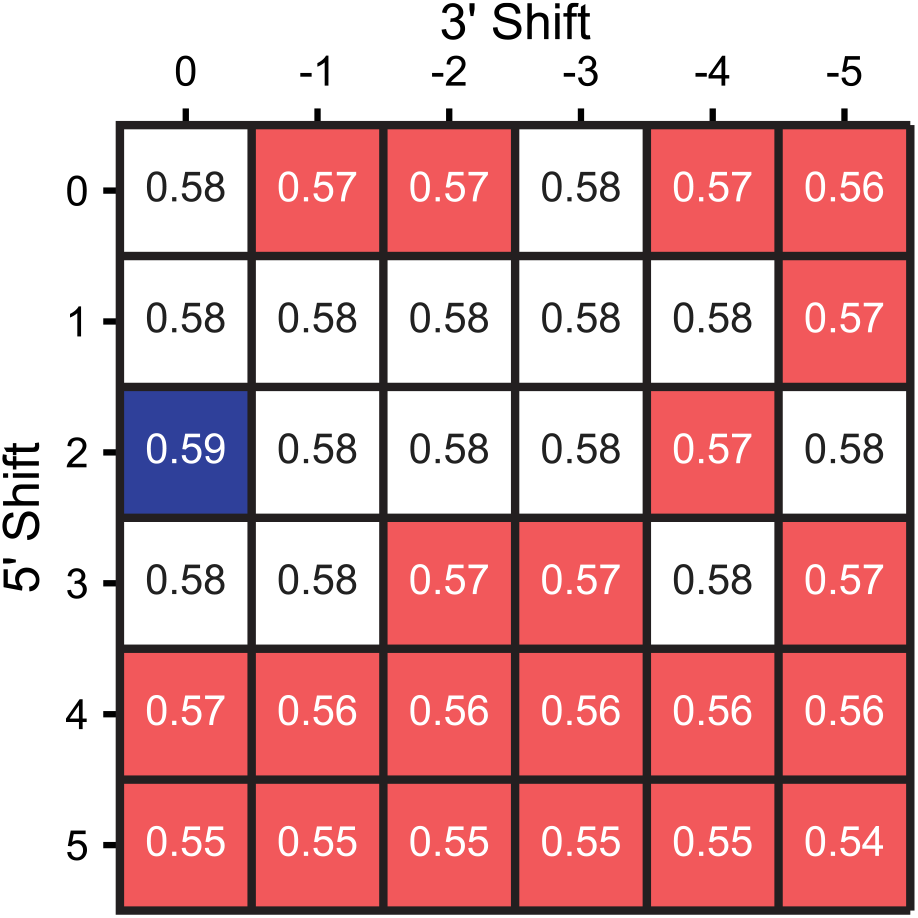
Effect of 5’ and 3’ shifts in site assignment on through-space distances. Identification of short distance interactions, as examined by receiver operating characteristic (ROC) curve analysis. Classifier: Inter-nucleotide distance less than 15 Å with a contact distance > 10 (see Methods), based on normalized deletion rate. Mean area under curve (AUC) values for a set of SHAPE-JuMP experiments, performed using five model RNAs (see Methods), as a function of 5’ or 3’ shift, are shown. Red, white, and blue coloring indicate AUC below, at, or above mean AUC value.

**Figure S5:**
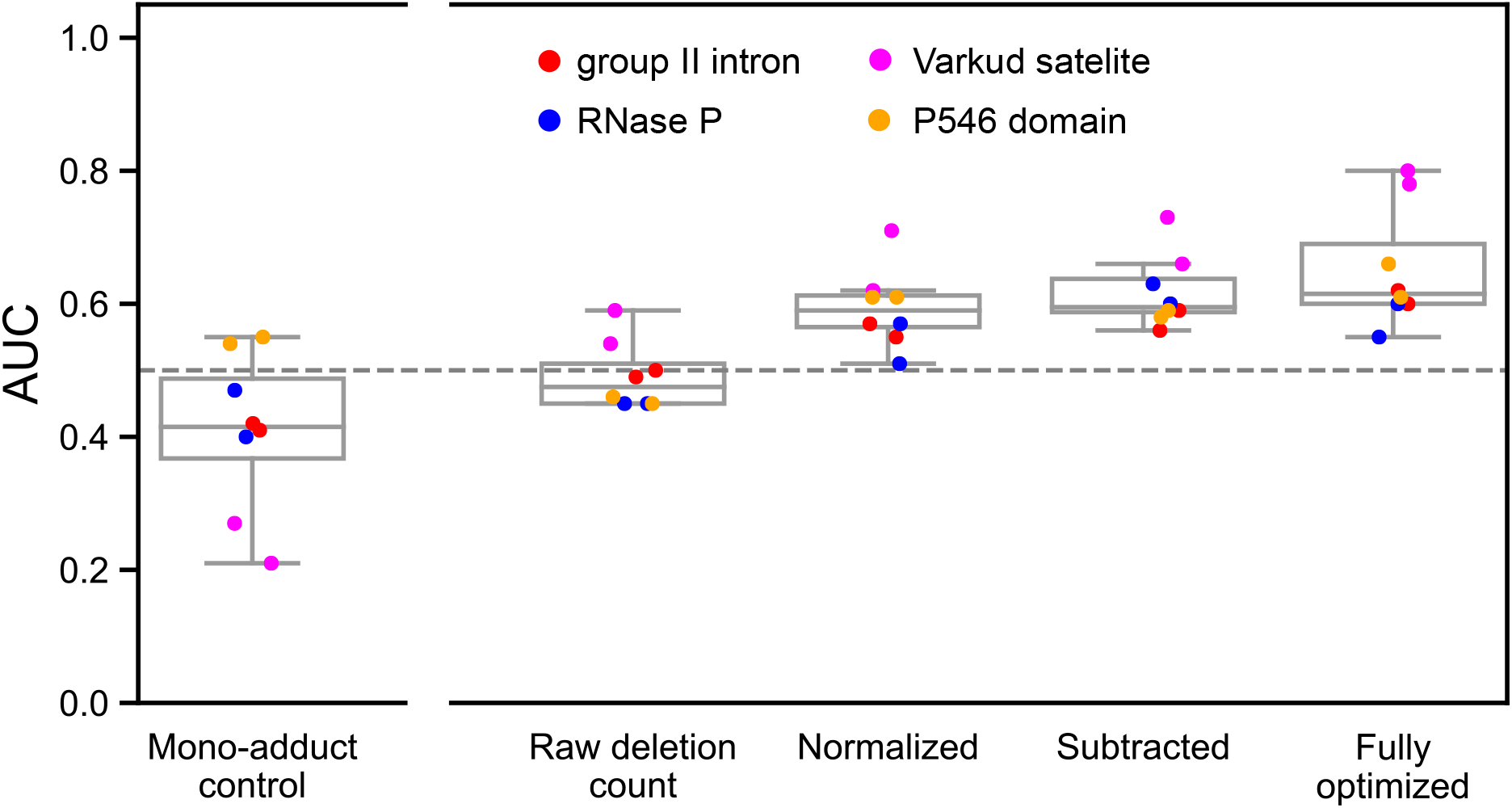
Progress of ShapeJumper optimization steps for through-space interaction identification. AUC values summarize the results of replicate experiments in terms of ability to measure close-in-space interactions, defined as through-space distances less than 15 Å and contact distances greater than 10. Each column represents a step in the ShapeJumper pipeline. The mono-adduct control shows the AUC for (non-crosslinked) IA samples after processing by the optimized pipeline.

